# Chemical unfolding of protein domains induces shape change in programmed protein hydrogels

**DOI:** 10.1101/711325

**Authors:** Luai R. Khoury, Ionel Popa

**Affiliations:** Department of Physics, University of Wisconsin-Milwaukee, 3135 North Maryland Ave., Milwaukee, Wisconsin 53211, United States

**Keywords:** programmable biomaterials, protein-based hydrogels, shape-memory hydrogels, protein folding mechanics

## Abstract

Programmable behavior combined with tailored stiffness and tunable biomechanical response are key requirements for developing successful materials. However, these properties are still an elusive goal for protein-based biomaterials. Here, we present a new method based on protein-polymer interactions, to manipulate the stiffness of protein-based hydrogels made from bovine serum albumin (BSA) by using polyelectrolytes such as poly(Ethelene)imine (PEI) and poly-L-lysine (PLL) at various concentrations. This approach confers protein-hydrogels tunable wide-range stiffness, from ~ 10 - 60 kPa when treated with PEI, without affecting the protein mechanics and nanostructure. We ascribe the increase in stiffness to the synergistic effect of the non-covalent electrostatic polymer-protein interaction, as well as the polymer-shell that stabilizes the protein domains nanomechanics. We use the 6-fold increase in stiffness induced by PEI to program BSA-hydrogels in various shapes. By utilizing the characteristic protein unfolding we can induce reversible shape-memory behavior of these composite materials using chemical denaturing solutions. We anticipate this novel approach based on protein engineering and polymer reinforcing will enable the development and investigation of new smart biomaterials and extend protein hydrogel capabilities beyond their conventional applications.

---

Polymer – based hydrogels have found broad applications in tissue engineering, drug delivery, soft robotics and actuators^1,2^, and their viscoelasticity can drive stem cell fate and activity^3^. However, these hydrogels possessed limited mechanical strength, are prone to permanent breakage, lack dynamic switches and reversible shape^2^. Several approaches have been proposed to improve their stiffness and extensibility. One method is through using a double-network crosslinking strategy, either by secondary polymer network, using multivalent ions, or by nanoparticles^4–10^. Furthermore, polymer hydrogels are being used in shape-memory field. For example, thermoplastic polymers-based hydrogels can display shape memory response as a function of temperature, which is of great importance to soft robotics and biomedical applications^11^. Additionally, supramolecular interactions between chains based on reversible hydrogen bonds, metal-coordination or dynamic covalent bonds have been recently introduced to improve hydrogels functionality, but in all these examples the structure of the primary network changes during shape-morphing cycles^8,12,13^.

In the last decade, protein hydrogels based on globular proteins were proposed as a novel biomaterial that may have a wide use in biomedical applications and research^14^. These hydrogels are intrinsically biocompatible, biologically diverse, and can use the unfolding response or tertiary structure for energy storage and release ^10,15–17^. Currently the stiffness of protein-based hydrogels has a narrow tunability range, limited at the lower end by the minimum protein concentration required for gelation, and at the higher end by the solubility of the protein. 10,16. It has been challenging to obtain the same smart behavior as that of polymer-based hydrogels, in part due to the limited range where solvents, temperatures and concentrations can be used. Proteins generally require water-based solvents, a narrow salt and pH range, and the working temperature to obtain biomaterials cannot exceed values well above 37°C. Furthermore, the range of concentrations that can be used to obtain hydrogels is narrow ^14,16,18^. At the lower end, a too low protein concentration leads to incomplete network formation, and soft gels with irreversible deformations under strain, due to incomplete cross-linking. At the upper end, while the final stiffness of protein-hydrogels can be improved with increasing protein concentration, and hence the cross-linking density, a major limitation comes from the maximum protein solubility. For example, hydrogels made from protein G, domain B1 (GB1), from SH3 or from chimera GB1-HP67 had a minimum gelation concentration of ~150 mg/mL and reached their solubility limit at ~180 mg/mL (which corresponds to ~1.3 to 3.2 mM)^18^. In this concentration range, the gels had a narrow range of stiffness, of ±0.6 kPa to ±3 kPa. For BSA based hydrogels, concentrations below 1 mM produce gels showing plastic deformation under force, while the maximum solubility of BSA (~4 mM) only yields hydrogels with stiffness of ~15 kPa, setting the upper limit achievable with this method^15^. An increase in the stiffness range for protein hydrogels would not only expand their applications, but also allow for shape programmable behavior. Such a shape-memory approach based on protein (unfolding) transitions does not currently exist.

Here, we report a method of producing hybrid protein-polymer hydrogels which have covalently-crosslinked protein network reinforced with physically adsorbed polyelectrolytes. We use a custom-made force rheometer which utilizes an analogue feed-back to expose protein-based hydrogels to various force protocols ^15,19^. We characterize the intake of various polyelectrolytes and determine the change in stiffness and folding of BSA based hydrogels. This approach provides the ability to fine-tune the stiffness of BSA-based hydrogel up to 6-fold without affecting the unfolding nanomechanics of proteins domains. Using this unique interaction between BSA and polymers, in combination with the unfolding response in chemical denaturants, we formulate protein-based hydrogels to display reversible shape-memory behavior.

## 1. Results

### 1.1. Polyelectrolytes can stiffen protein-based hydrogels

BSA is one of the most inexpensive and abundant proteins available. It has a negative charged at pH~7.4 due to several negatively charged amino acids patches distributed on its surface^20^. Our first goal was to determine the appropriate polyelectrolytes that can adsorb on BSA domains on the hydrogel matrix. First, hydrogels were synthesized inside semi-transparent tubes (inner diameter 558.8 μm) using 2 mM BSA and a photo-activated cross-linking reaction, which results in formation of covalent carbon-carbon bonds between two adjacent BSA domains via exposed tyrosine sites^15,19,21,22^. As previously reported, these gels are completely cross-linked and show completely reversible stress-relaxation behavior under force^15^. Following gelation, the BSA hydrogels were equilibrated in TRIS buffer (Tris 20 mM, NaCl 150 mM, pH~7.4) for 30 minutes at room temperature, then moved to one of the following polymer solutions: branched-polyethyleneimine (PEI) 10 kDa, poly-(L)-lysine (PLL) 10 kDa, and polyethylene glycol (PEG) 8 kDa, which were dissolved in the same TRIS buffer (Figure 1A). Following incubation for another 30 min in one of a specific polymer concentration at room temperature, the BSA hydrogels were stored back in TRIS buffer, to wash any unbounded polymer molecules from the treated samples. The hydrogels were then characterized using force-clamp rheometry^15^, scanning electron microscopy (SEM), and swelling ratio measurements.

**Figure 1.**
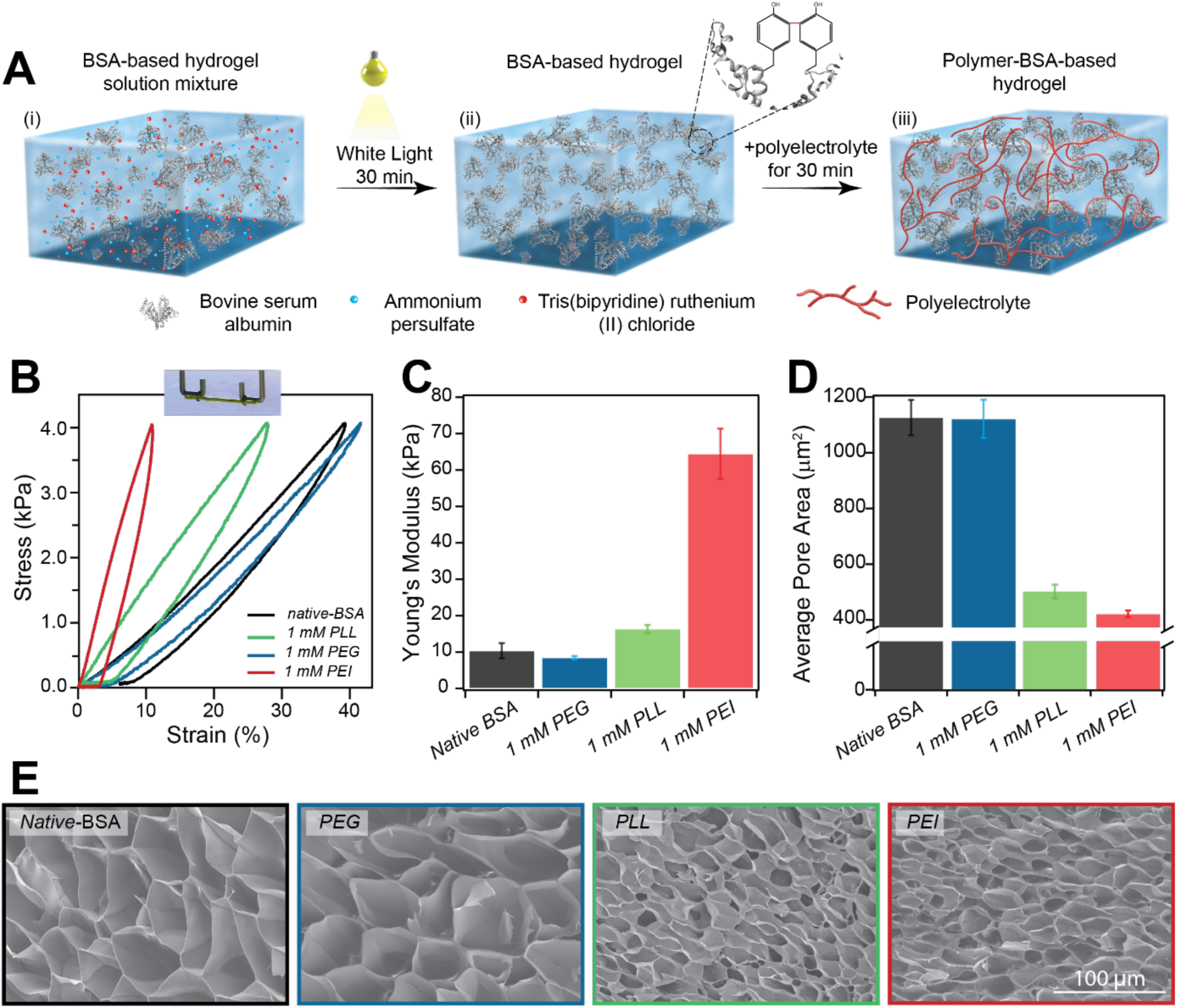
Synthesis of polyelectrolytes enforced protein-based hydrogels. (A) Strengthening process of BSA-based hydrogel using polymer-protein interaction: (i) BSA, ammonium persulfate (APS), and Tris(bipyridine) ruthenium (II) chloride ([(Ru(bpy)_3_]^2+^) are mixed together. (ii) the hydrogel mixture is exposed to white light for 30 minutes at room temperature (RT) which leads to a covalent cross-linking between BSA domains via two adjacent-exposed tyrosines amino acids (inset). Afterwards, the hydrogel is extruded into the TRIS solution. (iii) The TRIS solution is replaced with one of three polymer solutions for 30 minutes: PEI, PLL, or PEG. Thereafter, the hydrogel is immersed into the polyelectrolyte solution for 30 minutes at RT, then moved to TRIS solution to remove any unbounded polymer molecules. (B) Stress-strain curves of native-BSA (black) and after incubation with 1 mM PEG (blue), PLL (green) and PEI (red). Inset: A scheme shows the tethered hydrogels to the FC rheometer hooks. (C) Average Young’s moduli calculated from stress-strain curves of *native-*BSA when treated 1 mM of PEG (blue), PLL (green) and PEI (red). (D) Average pore-size values of native-BSA (black) and after incubation with 1 mM PEG (blue), PLL (green) and PEI (red) samples as derived from SEM images analysis. The error bars represent the standard deviation of the mean measurements. (E) SEM images of native-BSA and after incubation with the same polyelectrolytes.

The mechanical response of both native and polymer-treated hydrogels was measured using a force-ramp protocol where the stress was linearly increased and decreased with a rate of 40 Pa/s. Using this approach, we determine the Young’s modulus from the initial slope of each stress-strain curve, which reports on the gel stiffness. Furthermore, as proteins unfold and refold at vastly different forces ^23^, the stress-strain curves show important hysteresis, which reports on the energy being dissipated due to these phase transitions^19,24^.

When the hydrogels are treated with a constant 1 mM polymer concentration, PEI had the largest effect on the measured stiffness, which increased from 10.3 ± 2 to 64.4 ± 6.8 kPa. PLL increased the Young’s modulus to 16.3 ± 1.1 kPa, while PEG did not induce any change (Figure 1B&C). This trend was mirrored by the hydrogel structure when characterized by SEM, where no significant change in pore-size was seen upon incubation with PEG (pore size 1126 ± 63 µm^2^ for native BSA, 1121 ± 68 µm^2^ for PEG). PLL and PEI induced a decrease in the pore size, with areas of 502 ± 24 µm^2^ and 421 ± 12 µm^2^, respectively (Figure 1D&E).

Our rheometry-based approach allowed us to easily asses the change in stiffness of BSA hydrogels treated with different concentrations of polyelectrolytes (Figure 2). PEG-treated hydrogels, where the polymer was in 0.25 mM to 3 mM range, did not show any increase in the Young’s modulus (Figure 2A(iii)). On the other hand, both PLL and PEI resulted in the stiffening of the exposed BSA hydrogel (Figure 2A(i) and Figure 2A(ii)). The effect is more pronounced with PEI, where we measure up to 6-fold increase in the gel stiffness (Figure 2B). While both polymers are positively charged at pH~7.4 and with similar molecular weight, PEI is branched and has a higher charge density and interacts more efficiently with negatively charged amino acid patches on the BSA surface. The increase in stiffness is mirrored also by the decrease in the hysteresis measured from stress-curve for each polymer concentration (Figure 2C). Additionally, measurement of the pore-size for each treated hydrogel sample using SEM showed that there is a shrinking trend in the pore size from 1126 ± 63 µm^2^ to 359 ± 21 µm^2^ with increasing PEI concentration, which correlates with the slight decrease in water content (Figure 2D-F). Interestingly, the wall thickness of the pores showed a positive correlation when increasing the PEI concertation up to 0.75 mM but decreased for PEI concentrations greater than 1 mM (Figure 2G).

**Figure 2.**
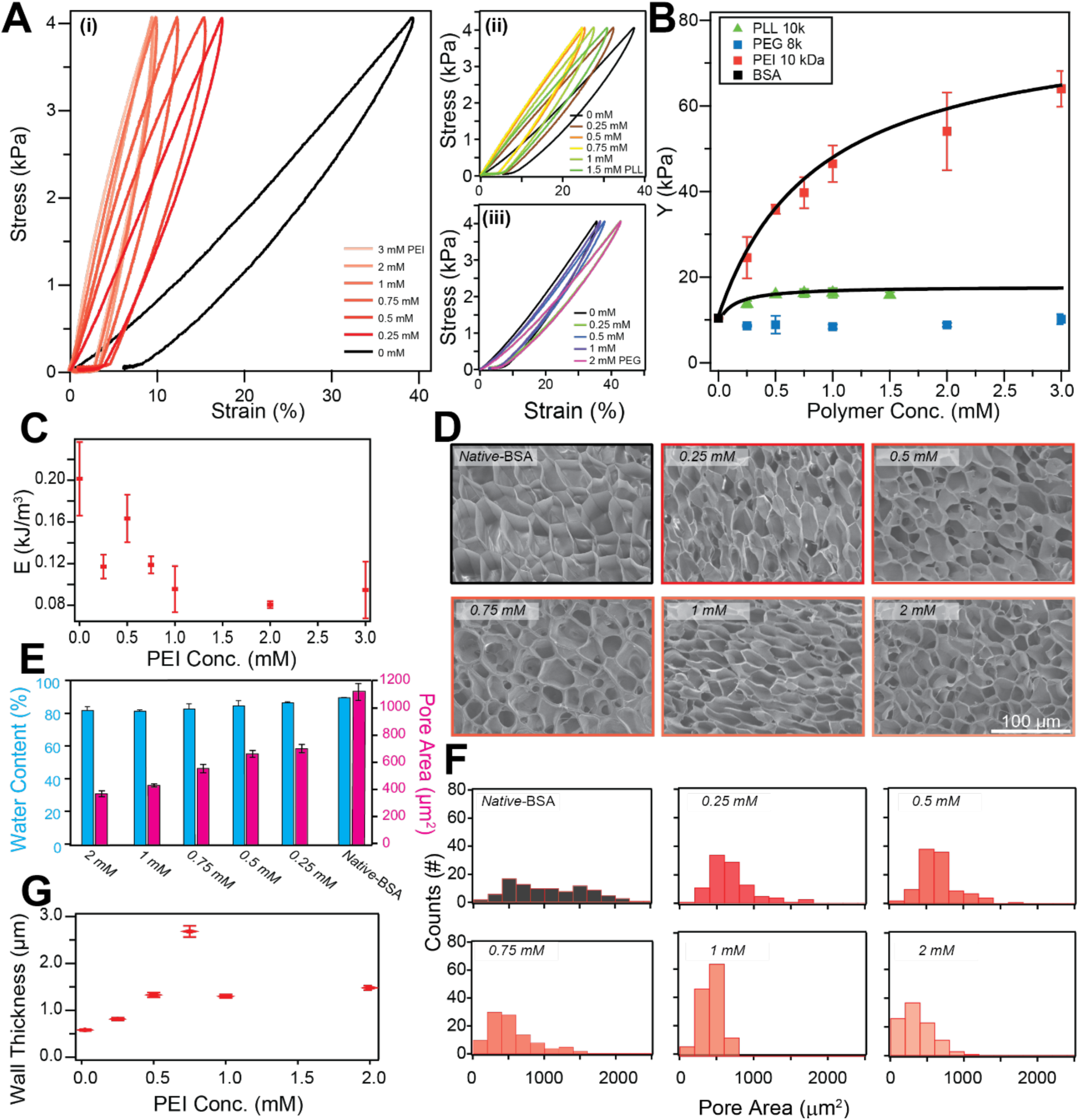
Characterizations of BSA hydrogels as a function of polyelectrolyte intake. (A) (i) Stress-strain curves of *native-*BSA and incubated with various concentrations of PEI ranging from 0 to 3 mM. (ii) Stress-strain curves of *native-*BSA and incubated with various concentrations of PLL ranging from 0 to 1.5 mM (iii) Stress-Strain curves of different BSA-based hydrogels treated with various PEG concentrations ranging from 0 to 2 mM. Each curve represents the average of three different measurements. (B) Average Young’s moduli calculated from stress-strain curves of *native-*BSA, and incubated with PEG (blue), PLL (green) and PEI (red) as a function of the polymer concentrations. Fits represent a Langmuir-like behavior, with equilibrium constants of *K*_*PLL*_ = 1.1 and *K*_*PEI*_ = 5.3 Mm^−1^. (C) Relationship between PEI concentration and energy dissipation of treated BSA hydrogels. The energy dissipation was calculated from the hysteresis area enclosed in the stress–strain curves. (D) SEM images of BSA-based hydrogel samples incubated with PEI at various concentrations (E) Average pore-area size and water content measurements of native and PEI treated hydrogel samples. The error bars represent the standard deviation of at least three different measurements. (F) Pore-area size distribution histograms as derived from SEM images. (G) The relationship between will thickness of the pores and PEI concentration.

### 1.2. The synergistic effect of polymer-protein interactions on tuning BSA-based hydrogel stiffness

To decouple the response of protein (un)folding mechanics from the intrinsic elasticity coming from the polymer-protein interaction, we used Guanidinium Hydrochloride (GuHCl) 6 M, which acts as chemical denaturant^19^. Addition of the GuHCl on native BSA hydrogels manifests in two ways: it softens the gels and removes the hysteresis^15,25^. The softening comes from the fact that folded linked proteins are ~20x stiffer than unfolded polypeptide chains^26^. The disappearance of the hysteresis in stress-strain curves is a benchmark for the lack of tertiary and secondary structure, here induced through chemical denaturation (Figure 3A and B). Interestingly, when adding GuHCl to protein hydrogels treated with PEI, the hysteresis disappears as expected, but the stiffness is higher than that of the native BSA gels in TRIS (~ 18 kPa vs **~** 10 kPa, Figure 3A and B). After washing out the GuHCl salts from the PEI-treated hydrogel sample by immersing it in TRIS solution, the BSA domains refold back to their native state and the hydrogel regains its initial stiffness (~ 60 kPa) and shows a similar hysteresis as before the immersion in the GuHCl solution (Figure3 A and B). Mechanical unfolding of protein domains enables BSA hydrogels, with or without PEI treatment, to maintain their elastic behavior all the way to the breaking force. However, due to its stiffening effect, PEI increased the maximum stress that a BSA hydrogel can sustain (Figure 3C).

**Figure 3.**
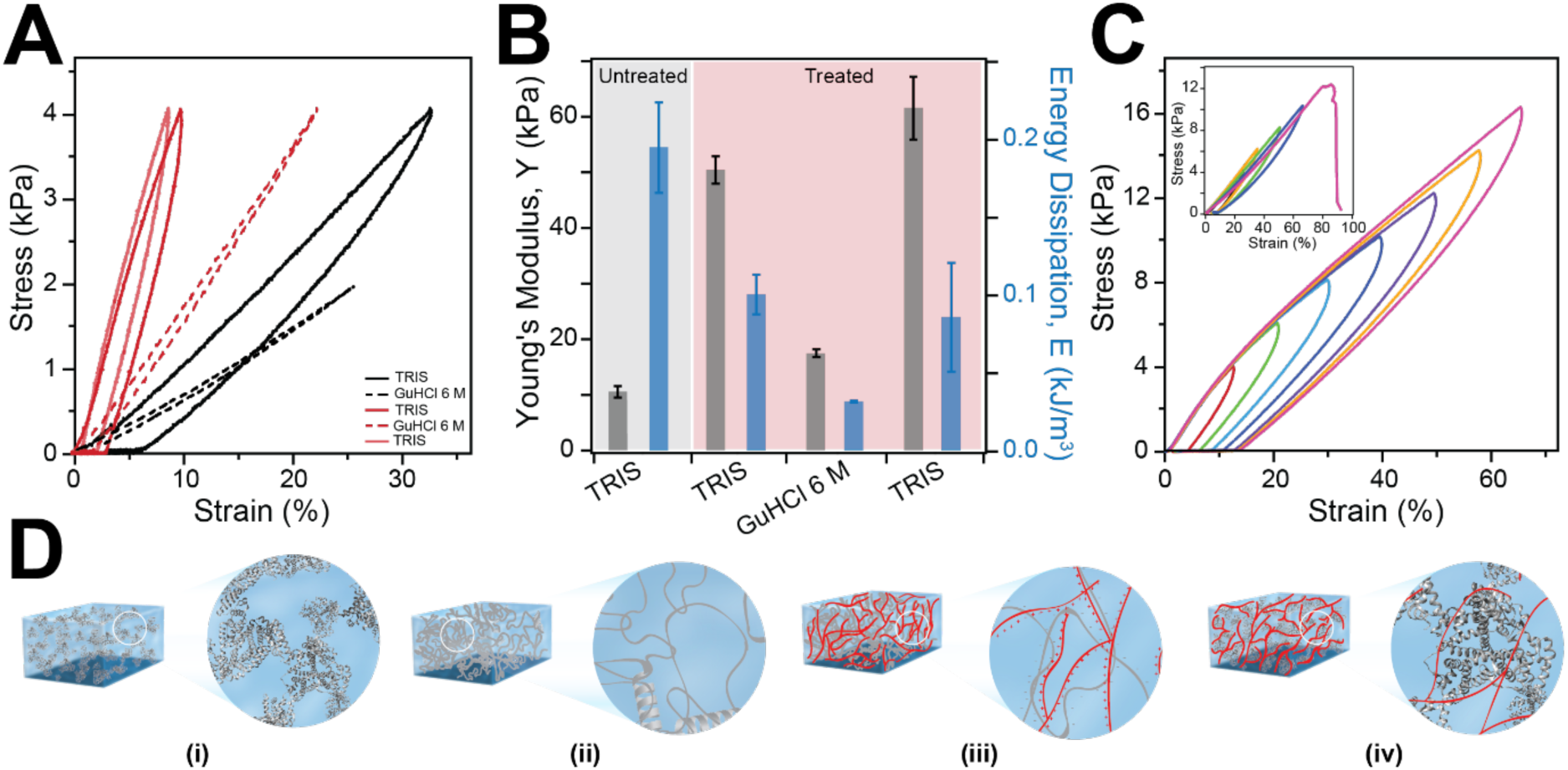
PEI stiffens BSA-hydrogels without preventing chemical denaturation of protein domains. (A) Stress-strain curves of same BSA-based hydrogel sample in TRIS and 6 M GuHCl before and after treatment with 2 mM PEI solution. This reversible behavior opens the door for shape memory and shape change in chemical denaturants. (B) Average Young’s moduli and energy dissipation of same BSA-based hydrogel sample in TRIS and 6 M GuHCl before and after treatment with 2 mM PEI solution. (C) Stress-strain of PEI-treated BSA-based hydrogel sample showing that it can withstand successive loading-unloading cycles with increasing final stresses while showing a pronounced hysteresis and recovery. Inset: Stress-strain of BSA-based hydrogel sample failed after successive loading-unloading cycles of increasing force. (D) Proposed mechanisms for the strengthening of treated PEI-BSA-based hydrogel: (i) Covalent crosslinking between BSA domains. (ii) Protein (un)folding nanomechanics. (iii) PEI molecules stabilize a BSA domain by forming a protective polymeric shell, and (iv) Non-covalent electrostatic crosslinking between BSA domains and PEI molecules.

### 1.3. Programming and manipulating shape-memory of BSA-based hydrogel using BSA-PEI interactions via denaturant solution

The PEI-BSA interactions inside the hydrogel matrix provide a new vista for constructing moldable and shape-memory biomaterials based on globular proteins. Above, we observed that PEI can strengthen the BSA-based hydrogel sample and exhibited a good recovery (Figure 3A-C). We used this phenomenon to program BSA-based hydrogels in various shapes. We demonstrate this approach using different 3D-printed molds (Figure 4A). A BSA-based hydrogel was immobilized to obtain a W- and spring-like shapes, and programmed by immersing it in a 2 mM PEI solution for 30 min at room temperature. Following a wash step with TRIS buffer, the gel preserved its programmed shape in TRIS buffer (Figure 4B). To disrupt the programmed shape of the BSA-based hydrogel, we used 6 M GuHCl denaturant solution. The denaturating solution triggers protein unfolding (Figure 4B and Figure S3) that results in the macroscopic loss of the programmed shape (Figure 4B). When the molded sample is moved back into TRIS, the gel regains its PEI-programmed shape, after a couple of minutes (Figure 4B and Movie S1).

**Figure 4.**
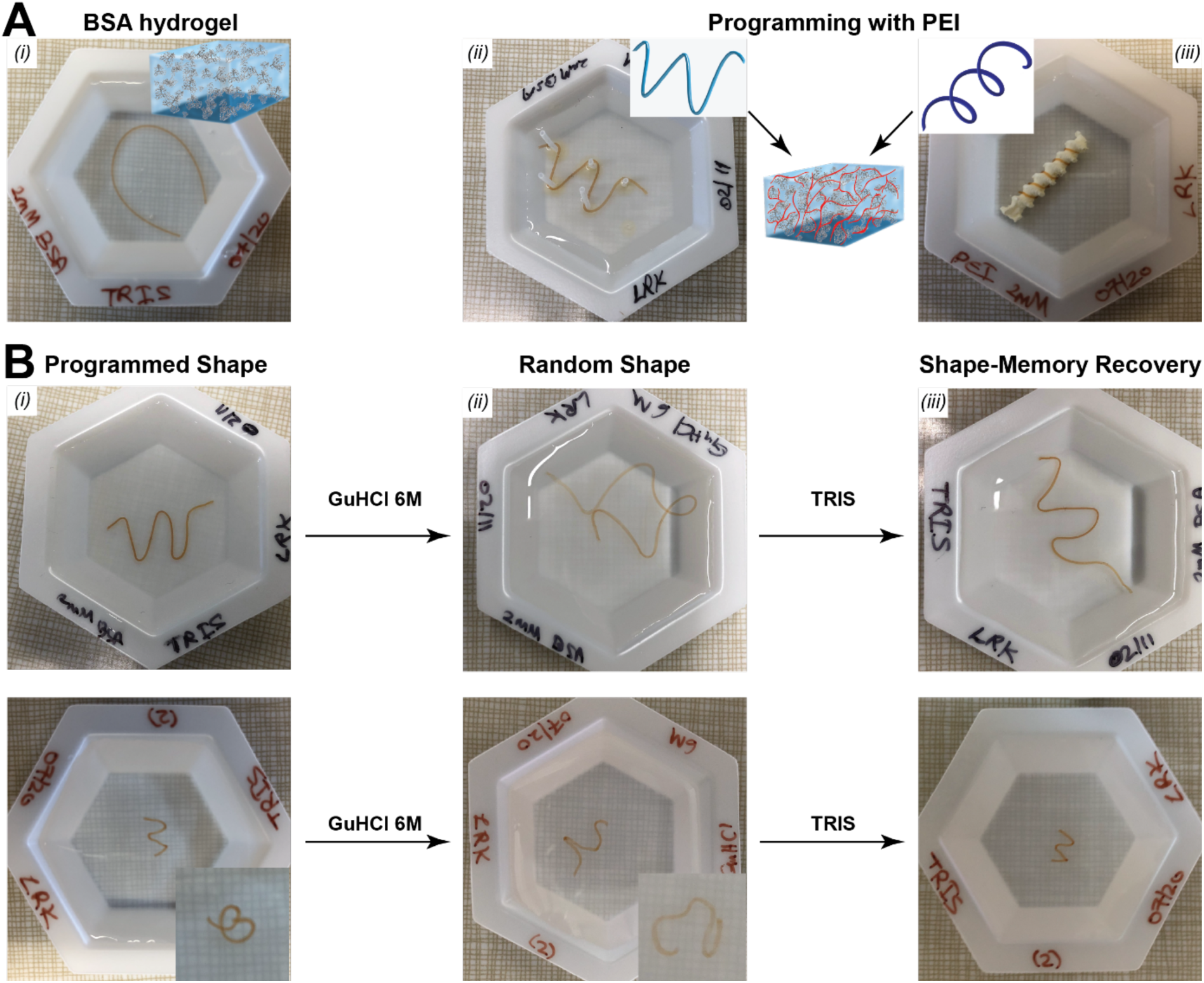
Shape*-*memory recovery process of BSA-2mM PEI based hydrogel with various shapes. (A) (i) 2 mM BSA-based hydrogel extruded from PTFE tube into TRIS buffer; (ii) A 2 mM BSA-based hydrogels programmed into W-like by treating with 2 mM PEI solution. (iii) A 2 mM BSA-based hydrogel programmed into a spring-like shape, using a 3D-printed screw-like shape structure, by immersing it in 2 mM PEI solution. (B) The programmed shapes of the PEI -treated BSA-based hydrogels in TRIS solution to washout any unbounded PEI molecules. The hydrogels retained their programmed shapes. Inset (bottom): Close-up of the programmed BSA-based hydrogel sample from a different angle. Then, the programmed BSA-based hydrogel samples moved into 6 M GuHCl denaturant solution that causes unfolding of the BSA domains and leads to a random temporary shape of the hydrogel samples. Inset (bottom): Close-up of a random-shape BSA-based hydrogel inside a 6 M GuHCl solution from different angle. Thereafter, the hydrogels samples recovered their programmed shapes after moving them into the TRIS solution. The GuHCl salt were being washed out and the BSA domains refold leading to hydrogel initial shape recovery in a couple of minutes.

## 3. Discussion

For protein hydrogels made from structured proteins, a unique viscoelastic property comes from the nanoscopic response to force of their constituent domains, which can reversibly unfold and extend 4 to 10 times their initial folded length^14,26^. This unique response to force, combined with their biocompatibility and diverse functional spectrum, place protein hydrogels at the forefront of bioengineering. However, they still have major drawbacks. (i) Protein hydrogels typically show weak mechanical integrity and increasing the number of cross-linking sites can improve their stiffness, but at the expense of a narrower tunability^14,27^. ^18^For BSA, the minimum gelation concentration is ~0.7 mM, while the saturation concentration is ~4 mM, which translated into a Young’s moduli range between 2.5 to 15 kPa^15^. When treated with PEI, BSA-based hydrogels (2 mM) showed a significant increasein the Young’s modulus, up to ~65 kPa (~ 6-fold increase), and a wide range of stiffness tunability, ranging from 10 to 60 kPa (Figure 2). (ii) Hydrogels also tend to be weak and brittle, with low dissipation energy and elastic moduli^28^. Several different approaches were proposed for polymer-based hydrogels to circumvent this important flaw, using double^7^ or triple^29^ overlapping networks. Typically, the structural failure of the first network is mitigated by the take-over of the secondary network. Here too, by immersing BSA hydrogels in polyelectrolytes such as PEI, we generate a double-network. However, polymer-treated protein hydrogels operate on a different mechanism, where the primary network can dissipate energy through protein (un)folding nanomechanics, and the secondary polymer network acts to tune the stiffness and reinforce the hydrogel. In our case, PEI adsorption does not only serve as a secondary supportive network, but also contributes synergistically to the mechanical stability of the BSA domains inside the hydrogel matrix. As the concentration of PEI increases, the measured hysteresis in the stress-strain curves decays to a constant value at 0.75 mM of PEI (Figure 2C). Furthermore, since unfolding and refolding of protein domains is a reversible process, and a large amount of mechanical work can be dissipated during the protein (un)folding transitions, the native BSA gels do not show plastic permanent deformations until breaking, which occurs at ~11 kPa for PEI-free BSA hydrogels. Whereas, the PEI treatment increased the breaking force beyond 24 kPa, (Figure 3C). Furthermore, PEI enables reproducible behavior while the final force is increased successively, without impairing the backbone structure of the hydrogel, these behaviors were no even possible with untreated BSA-based hydrogels (Figure 3C).

The significant increase in stiffness, extensibility and recovery of BSA hydrogels treated with PEI can now allow us to program these hydrogels into a specific shape, which can enable these materials to extend their use into different applications such as soft robotics and actuators^12,13^. We accomplished this by mounting the BSA hydrogel on a mold and programmed its shape by immersing it in PEI solutions (Figure 4). To trigger shape changes, we used the unique response of proteins to chemical denaturants, such as GuHCl. PEI incubated BSA hydrogels display a change in stiffness from ~60 kPa in TRIS buffer to ~18 kPa and lose their hysteresis, as shown stress-strain curves (Figure 3A). This significant change in stiffness in chemical denaturant leads to loss of the programmed shape, as the BSA domains forming the skeleton are denatured. Importantly, this unfolding process is reversible, as the BSA gels recover both their initial Young’s modulus and energy dissipation behavior when immersed back from GuHCl to TRIS (Figure 3A-B). Macroscopically, this results in the recovery of the programmed shape during the washout of the GuHCl salts (Movie-S1). This approach demonstrates that incubation of polyelectrolyte with protein hydrogels does not only increase the attainable stiffness and tunability, but also allow hydrogels to operate in a stimuli-responsive manner. Other systems based on fibrillar proteins such as collagen use swelling and deswelling to actuate macroscopic movements^30^. Our system is unique, as it is able to reversibly switch macroscopically between extended and programmed shape in response to the nanoscopic unfolding and refolding of protein domains.

Finally, the experiments performed here allow us to also understand the synergistic strengthening mechanism. First, covalent cross-linking of BSA molecules at the tyrosine sites produce a network that responds to force in a fully reversible way in the sampled force range (Figure 3D(i)). Second, the (un)folding nanomechanics of BSA domains inside the hydrogel matrix change the elasticity and allow for large amounts of energy dissipation before physical damage of the network can occur (Figure 3D(ii)). Third, the non-covalently attached polyelectrolytes can form and break local bonds, allowing the gels to heal any structural damage inside the BSA network caused by the applied stress or strain (Figure 3D(iii)). Forth, the synergistic effect of non-covalent electrostatic polymer-protein interaction as well as the that the PEI molecule stabilizes the BSA proteins, acting as a shell around the folded domains without affecting BSA (un)folding nanomechanics (Figure 3B and 3D(iv)).

## 4. Conclusions

In summary, we have demonstrated a simple method to program the shape of protein hydrogels using polyelectrolytes and to induce a reversible shape-change using the unfolding-refolding response via chemical denaturants. This programming is possible due to the strengthening effect that PEI has on BSA hydrogels, which can change the Young’s modulus up to 6-fold its original value. The unique polymer-protein interaction inside the hydrogel matrix introduces electrolyte-treated protein hydrogels as a new player to the shape-memory field. While the reversible response is induced here by chemical denaturants, we anticipate that other protein (un)folding specific triggers, such as pH, light and temperature via nanoparticles or external triggers could be introduced in the future. Given the recent developments in designing heteropolymers and the huge library of proteins, it will be possible to generate new smart protein-based hydrogel biomaterials for further use as drug delivery vehicles ^31^, tissue engineering scaffolds^32^ smart actuators as artificial muscles, and soft robotics for delicate bio-applications^12,13^, limited largely only by imagination.

## Supporting Information

Supporting Information is available in SI file

## Acknowledgments

We thank Dr. Heather Owen for access to the Electron Microscopy Facility and help in acquiring scanning electron microscopy images. This research was funded, the Greater Milwaukee Foundation (Shaw Award), the University of Wisconsin-Milwaukee, and by the National Science Foundation (grant number MCB-1846143).

## Notes

The authors declare no conflict of interest.

